# Redox-dependent Structure and Dynamics of Macrophage Migration Inhibitory Factor Reveal Sites of Latent Allostery

**DOI:** 10.1101/2021.09.01.458630

**Authors:** Erin Skeens, Meagan Gadzuk-Shea, Dilip Shah, Vineet Bhandari, Devin K. Schweppe, Rebecca B. Berlow, George P. Lisi

**Author notes:** To whom correspondence should be addressed: George P. Lisi, Rebecca B. Berlow.

## Abstract

Macrophage migration inhibitory factor (MIF) is a multifunctional immunoregulatory protein that is a key player in the innate immune response. Given its overexpression at sites of inflammation in a wide range of diseases marked by increasingly oxidative cellular environment, a comprehensive structural understanding of how cellular redox conditions may impact the structure and function of MIF is necessary. We used solution NMR spectroscopy and mass spectrometry to investigate structural and dynamic signatures of MIF under varied solution redox conditions. Our results indicate that the MIF structure is modified and becomes increasingly dynamic in an oxidative environment, which may be a means to alter the MIF functional response in a redox-dependent manner. We identified latent allosteric sites within MIF that are redox-sensitive and mutational analysis reveals that loss of redox-responsive residues attenuates activation of the coreceptor CD74. Leveraging sites of redox-sensitivity therefore reveals an avenue to modulate MIF function in its “disease state” via structure-based drug design.

## Introduction

Macrophage migration inhibitory factor (MIF) is a ubiquitous human protein that plays a crucial role in the pathophysiology of inflammation.^1^ MIF is expressed by nearly every cell in the body^1^ and many of its effects are elicited by its binding to the chemokine receptors CXCR2 and CXCR4^2,3^ or the coreceptor CD74.^4-6^ Circulating levels are reported in the low μM range in healthy humans,^7-9^ but MIF is overexpressed under inflammatory conditions and has been linked to glucocorticoid overriding activity,^10^ asthma,^7,11^ rheumatoid arthritis,^12^ colitis,^13^ pancreatitis,^14^ acute respiratory distress syndrome (ARDS),^15^ and cancer^16-20^ as well as complications from COVID-19.^21^ Efforts to elucidate the pathogenic mechanism of MIF have attempted to connect its disease state to cellular redox properties,^11^ most recently by investigating its cysteine residues as conformational switches.^22^ Mutation of either C56 or C59 in a canonical ^56^CXXC^59^ (XX = AL in MIF) motif modulates the enzymatic and macrophage activation functions of MIF.^23,24^

An oxidative cellular environment is one of the hallmarks of inflammatory disease, characterized by excess reactive oxygen species (ROS) at sites of inflammation.^25^ Despite the acknowledged contribution of redox imbalance to asthma,^26,27^ ARDS,^28^ pulmonary fibrosis,^29^ and cancer,^30^ as well as the link between these pathologies and MIF, very little structural work has directly addressed the redox behavior of MIF. We wondered whether MIF could utilize redox imbalances to act as a pro-inflammatory sensor to toggle its structure based on the chemical environment of the cell. Altered conformations of MIF at local sites of inflammation could then modulate its function or create different modes of interaction with accessory proteins, thereby controlling downstream biological responses. In a first step toward exploring such a mechanism, we investigated the impact of solution redox conditions on the MIF structure and identified regions of the protein that are sensitive to oxidative environments. We used solution nuclear magnetic resonance (NMR) spectroscopy to assess the redox-dependent dynamics of MIF with spin relaxation experiments and used mass spectrometry to identify redox-modified cysteine residues. We then leveraged these findings, using further NMR structural studies and an *in vivo* assay, to confirm redox-sensitive amino acids as latent points of functional control in MIF.

## Results

### Oxidizing solution alters MIF structural stability

To assess the effect of solution redox potentials on the MIF structure, we performed far-UV circular dichroism experiments on redox-neutral MIF samples as well as MIF samples that had been oxidized or reduced. CD spectra of redox-altered MIF show characteristic α-β structure, with only minor differences (**Figure S1A**). These data are consistent with numerous MIF structures in the Protein Data Bank (PDB) that have no obvious alterations in the presence of substrates, inhibitors, or mutations. Despite similarly folded structures, temperature-dependent CD experiments (**Figure S1B**) show that an oxidizing environment destabilizes MIF to unfolding relative to reduced and redox-neutral MIF (Δ*T*_m_ = - 4.56 °C, ΔΔ*G* = - 2.65 kcal/mol, **Figure S1C**). This energetic difference (expressed per trimer) is modest, but suggests a loosening of the MIF structure that may affect its conformational ensemble.

### Redox-dependent NMR spectra of MIF highlight multiple conformations and altered dynamics

We used solution NMR to pinpoint regions of the MIF structure that are sensitive to the redox environment. ^1^H^15^N TROSY-HSQC spectral overlays of reduced, oxidized, and redox-neutral MIF (**Figure S2**) show that while reduced and redox-neutral samples are spectrally similar, more significant spectral changes are observed for MIF under oxidizing conditions. Redox-sensitive chemical shift perturbations (relative to redox-neutral MIF) are observed throughout the protein, including for residues at the N-terminal enzymatic active site, solvent channel, and monomer-monomer interface (**Figures 1A**). Additionally, for oxidized MIF samples, duplicate resonances are observed for 22 residues (**Figures 1B, S3**), suggesting that an oxidizing environment modulates an equilibrium between two conformations of MIF on the NMR timescale. Sites of slow exchange localize to the monomer-monomer interface (**Figure 1C**), indicating that the structure of this region of the protein is strongly affected by oxidizing solution, consistent with the lower thermal stability observed by CD. Perturbations to MIF are not exclusive to redox-active Cys residues, at times occurring 10 - 20 Å from the ^56^CALC^59^ motif or C80. However, Cys and Met residues serve as nucleation points for our observations, with a majority of redox-dependent changes to the MIF structure occurring proximal to these traditionally sensitive sites (**Figure S4**). These data are consistent with an alternate oxidized MIF conformation, “oxMIF,” most recently probed in a synthetic peptide comprising the MIF ^56^CALC^59^ sequence.^22^

**Figure 1.**
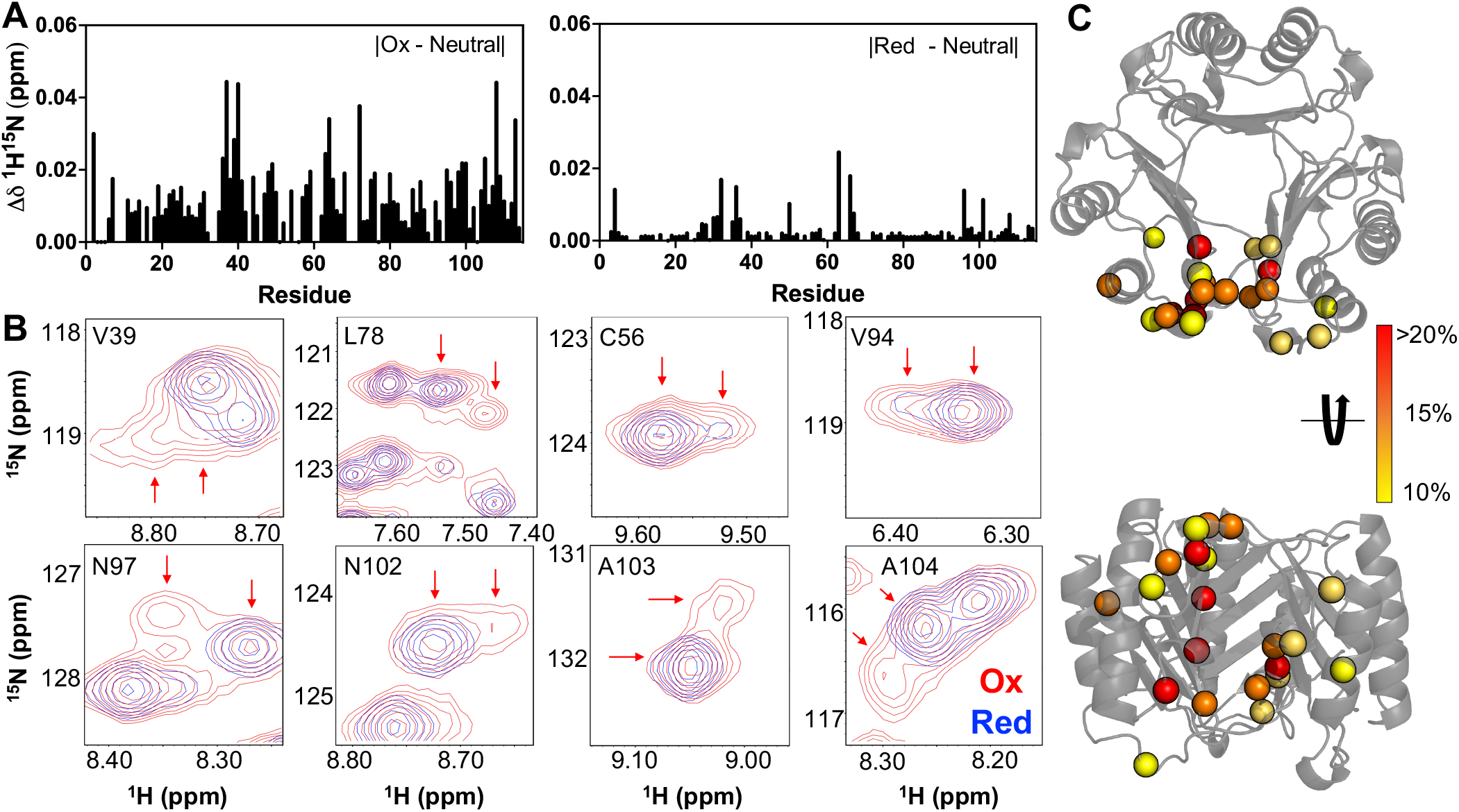
Redox-dependent changes in MIF detected by NMR. **(A)** ^1^H^15^N combined chemical shift perturbations (Δδ) for oxidized and reduced MIF, relative to a redox-neutral sample. **(B)** Selected resonances from ^1^H^15^N HSQC spectral overlays of reduced (blue) and oxidized (red) MIF. Sites of slow exchange observed under oxidizing conditions are indicated by arrows. Populations of MIF conformers determined from the volumes of slow exchange resonances are mapped onto the MIF trimer (PDB: 1MIF) in **(C)**, where the percentage indicated in the legend denotes the minor species (*i*.*e*. satellite peak). Volumes and populations of slow exchange resonances can be found in Figure S3.

Analysis of longitudinal (*R*_1_) and transverse (*R*_2_) NMR relaxation rates^31^ highlights local differences caused by redox conditions. Consistent with the altered thermodynamic parameters of the oxidized MIF samples, individual relaxation rates, as well as the *R*_1_*R*_2_ product, qualitatively imply that MIF exhibits heightened flexibility under oxidizing conditions (**Figure S5A**). Analysis of *R*_1_*R*_2_ values^31^ highlights 21 sites suggestive of μs – ms flexibility in oxMIF (*R*_1_*R*_2_ >1.5σ above the 10% trimmed mean of the data), 7 of which do not appear flexible in both redox-neutral and reduced MIF. Despite local patches of altered flexibility, the dynamic profiles of reduced and redox-neutral MIF are similar to each other. ^1^H-[^15^N] NOE values and order parameters also hint at patches of local flexibility in oxMIF, though fast timescale dynamics do not differ substantially between samples. Residues with relaxation parameters suggestive of μs – ms flexibility are mapped onto the MIF trimer and monomer structures in **Figure S5**. Correlation plots of *R*_1_*R*_2_ and *S*^2^ parameters for reduced and oxidized MIF reveal a weaker correlation for *R*_1_*R*_2_, suggesting that MIF experiences greater dynamic changes on the μs – ms timescale over a range of redox conditions (**Figure 2A**). Contributions to the variations in *R*_*1*_*R*_*2*_ parameters are most strongly affected by oxidizing solution conditions (**Figure 2B, left panel**), while reducing conditions contribute more significantly to the ps – ns dynamics of MIF via *S*^2^ (**Figure 2B, right panel**). Redox-sensitive residues with apparent μs – ms flexibility include T30, G50, L58, S63, I64*, K66, R73, L83, A84, R86, R93, I96, N97*, N109*, N110, T112, and A114, and those with apparent ps – ns fluctuations are F3, T7, L26, T30, D44, I64*, K66, K77, L83, L87, S90, Y95*, D100, A103, and S111* (**Figure 3C**). Asterisks indicate residues previously identified as controlling MIF enzymatic activity and CD74 activation via direct interaction or through allosteric signaling,^6,32,33^ indicating redox-dependent perturbations in MIF may have functional consequences.

**Figure 2.**
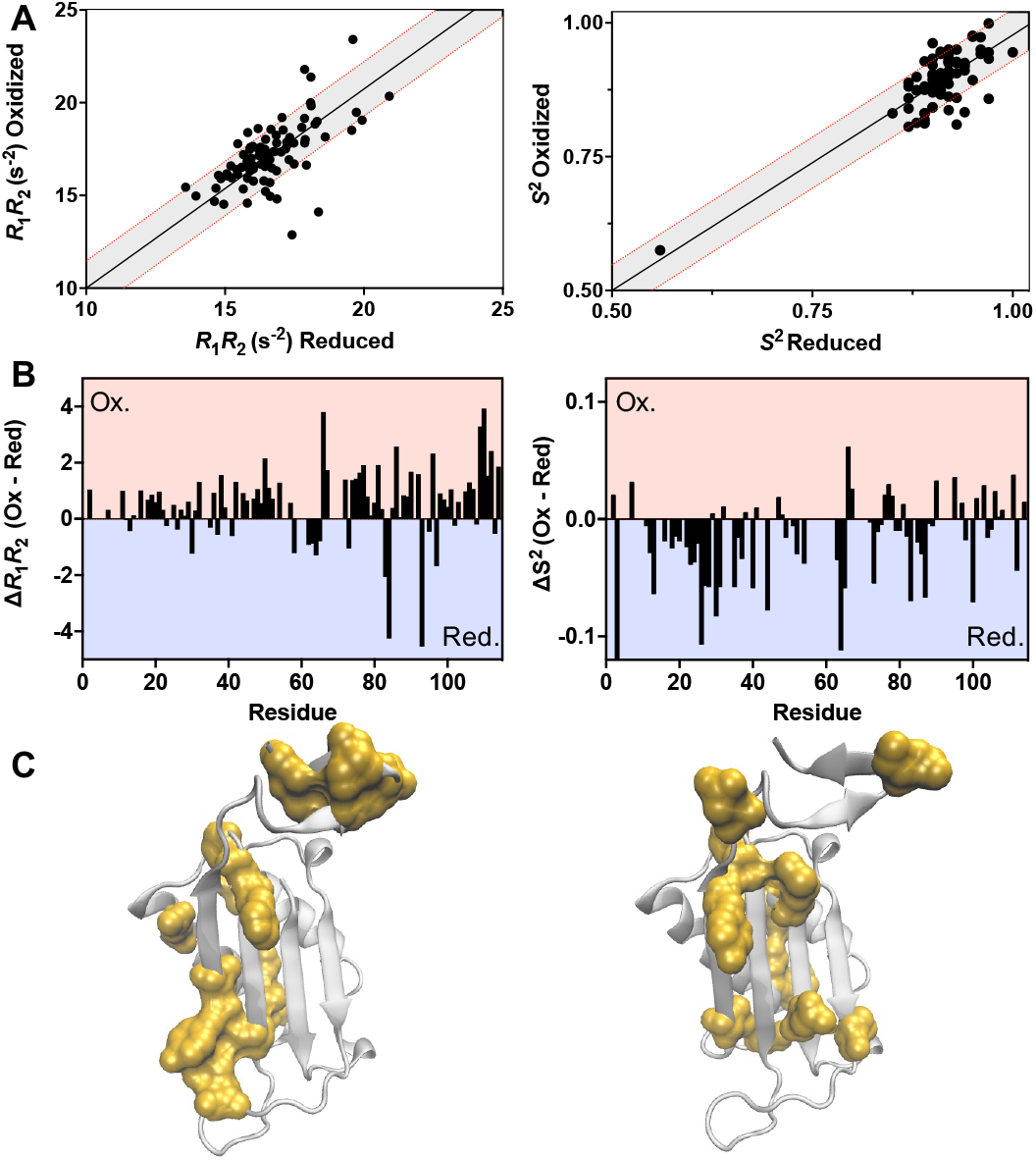
**(A)** Correlation plots of the *R*_1_*R*_2_ (left) and *S*^2^ (right) parameters for oxidized and reduced MIF. Gray shaded areas represent ±1.5σ from the linear least-squares fits of the data (black lines). **(B)** Per-residue differences in *R*_1_*R*_2_ (left) and *S*^2^ (right) plotted as oxidized – reduced. Bars in the shaded red area denote a greater value in oxMIF, while those in the blue area denote a greater value in reduced MIF. **(C)** Residues outside of the 1.5σ correlation boundaries in **(A)** are mapped onto the MIF monomer (PDB: 1MIF), highlighting regions of the protein with redox-dependent differences in μs – ms (left) and ps – ns (right) dynamics.

**Figure 3.**
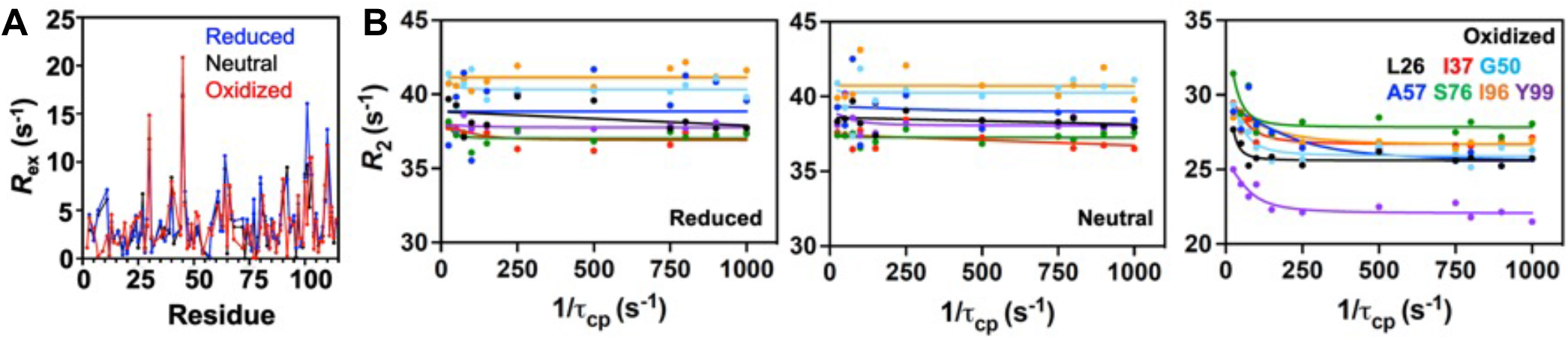
**(A)** Per-residue exchange contribution to relaxation (*R*_ex_) for reduced (blue), redox-neutral (black), and oxidized MIF (red). **(B)** CPMG relaxation dispersion profiles for selected MIF residues collected in reducing (left), redox-neutral (center), or oxidizing (right) solution. These sites display evidence of μs – ms flexibility under oxidizing conditions.

To confirm the influence of solution redox potential on μs – ms dynamics in MIF, we quantified the exchange contribution to transverse relaxation, *R*_ex_, via CPMG relaxation dispersion experiments. The average global *R*_ex_ value (<*R*_ex_>) for MIF is fairly similar across the redox regimes tested, with local differences arising primarily under oxidizing conditions (**Figure 3A**). Numerous sites of μs – ms dynamics are observed in all redox-modulated MIF samples (**Figure S6**), however, oxidizing solution stimulates the most widespread relaxation dispersion, affecting 39 residues (∼34 %) in the MIF structure. These data are consistent with *R*_*1*_*R*_*2*_ values that estimated a stronger contribution from oxidizing solution to μs – ms motions (**Figure 2**).

Regions of MIF with known functional importance become flexible, including the monomer interface (L26, I37, G50) and solvent channel allosteric site (I96, Y99). Despite similar dynamics in large portions of MIF, **Figure 3B** highlights selected sites that only appear flexible under oxidizing conditions. With the exception of the solvent channel, these flexible sites have not been well characterized for functional impact, suggesting that studies of MIF redox dependence can reveal new target residues for structure-function correlations.

### Cysteine 80 is a selectively modified redox switch

Prior work with a MIF epitope^22^ suggested that C80 functions as a redox molecular switch. However, redox sensitivity of C80 in the full-length trimeric protein is unknown. We performed quantitative mass spectrometry to explore the possibility that C80 could be selectively modified by redox reagents. Stepwise alkylation of MIF under each redox condition with iodoacetamide (IAA) and N-ethylmaleimide (NEM) yields critical insight into the oxidation state of C80 in the intact MIF trimer (**Figures 4, S7, S8**). Using NEM-bound C80 as a proxy for the abundance of cystine residues reveals an increase from 75.1% to 91.3% cystine abundance between redox-neutral and oxidizing conditions (**Figure 4B**). Under redox-neutral conditions, C80 exhibits the greatest variance in cystine abundance, further highlighting the reactivity and sensitivity of this residue to environmental conditions. The relative abundance of cystine is also sensitive to the amount of oxidative stress (**Figure S8A**). Increasing the concentration of oxidant in solution from 1:1 to 2:1 shifts the C80 cystine relative abundance from 83.1% to 91.3%. For comparison, **Figure S7B** shows the expected contributions to the total ion intensity of C80 modified with IAA or NEM if C80 is in a fully reduced, 50% reduced, or fully oxidized state. In addition to C80, MIF also contains two cysteine residues in its ^56^CALC^59^ motif; however, the peptide fragment containing C56 and C59 is a large 34-mer, precluding quantitation by the MS methods utilized here. Although we focus on C80, we cannot rule out a significant role for other cysteines in MIF redox dependence.

**Figure 4.**
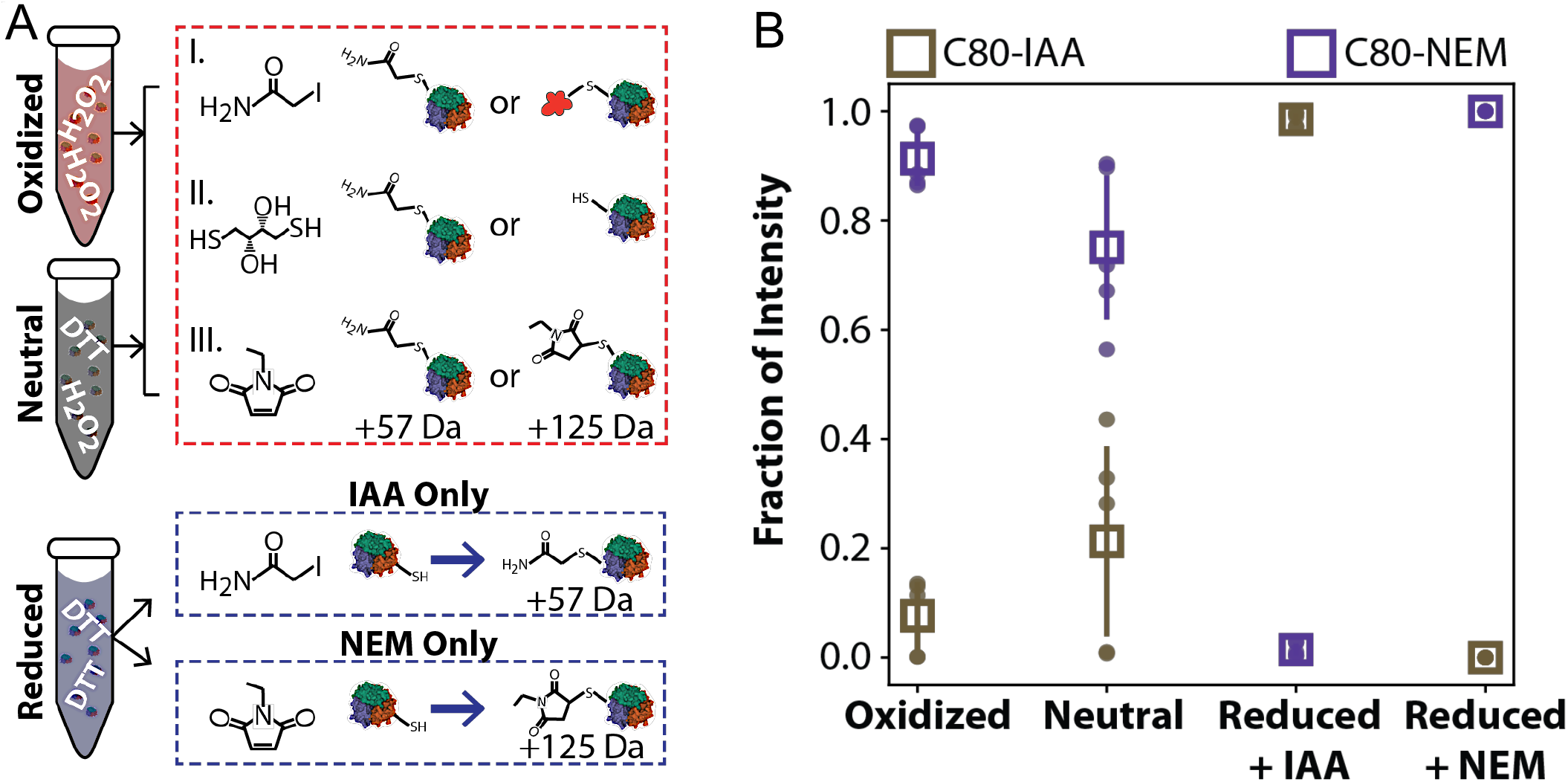
**(A)** Stepwise alkylation of MIF cysteine residues to determine the relative abundance of oxidized residues via mass spectrometry. The alkylation workflow involved three reaction steps (red, top panel). MIF equilibrated under reducing conditions was treated with only one alkylating agent and served as a quantitative control (blue, bottom panels). **(B)** Relative quantitation of C80 alkylation revealed conformational dependence and sensitivity to the surrounding redox environment.

### Mutation of redox-sensitive residues alters MIF structure and biological function

To test the possibility that the redox sensitivity of MIF could reveal sites of functional impact that are not otherwise obvious, we created single point mutations at two residues displaying redox-dependent structural and/or dynamic changes, lysine 66 (K66) and cysteine 80 (C80). K66 was found to have a large variation in the *R*_1_*R*_2_ parameter in oxMIF, as well as a low ^1^H-[^15^N] NOE in reduced MIF (Figures 2, S5), indicating that it undergoes significant changes in flexibility between the oxidized and reduced states. C80 also exhibits dynamic changes suggestive of redox-altered μs – ms chemical exchange and was shown by mass spectrometry to be selectively modified under oxidizing conditions (Figures 4, S5-8). ^1^H^15^N TROSY-HSQC NMR spectra of the K66A and C80A variants were collected in redox-neutral, reducing, and oxidizing solutions. Baseline structural changes caused by the K66A and C80A mutations are significant under redox-neutral conditions, particularly for the C80A variant, though CD spectroscopy indicates the variants are fully folded at NMR sample temperatures (Figures S9, S10). Interestingly, chemical shift perturbations observed for K66A MIF under oxidizing and reducing conditions are modest when compared to a redox-neutral K66A sample (Figures 5A, 5B). Most notably, oxidized K66A exhibits a chemical shift profile much smaller in magnitude than that observed in the corresponding wt-MIF experiments. In addition, reduced K66A exhibits larger chemical shift perturbations than oxidized in both number and magnitude, the inverse of what was found for wt-MIF. These data alone suggest that K66 may play a role in modulating the redox-dependent structural changes of wt-MIF, which are substantially diminished upon mutation. Likewise, the C80A variant strongly attenuates redox-sensitive NMR chemical shift changes in both reduced and oxMIF. These data, along with mass spectrometry, indicate that C80 modulates the conformational landscape of MIF under oxidative conditions.

**Figure 5.**
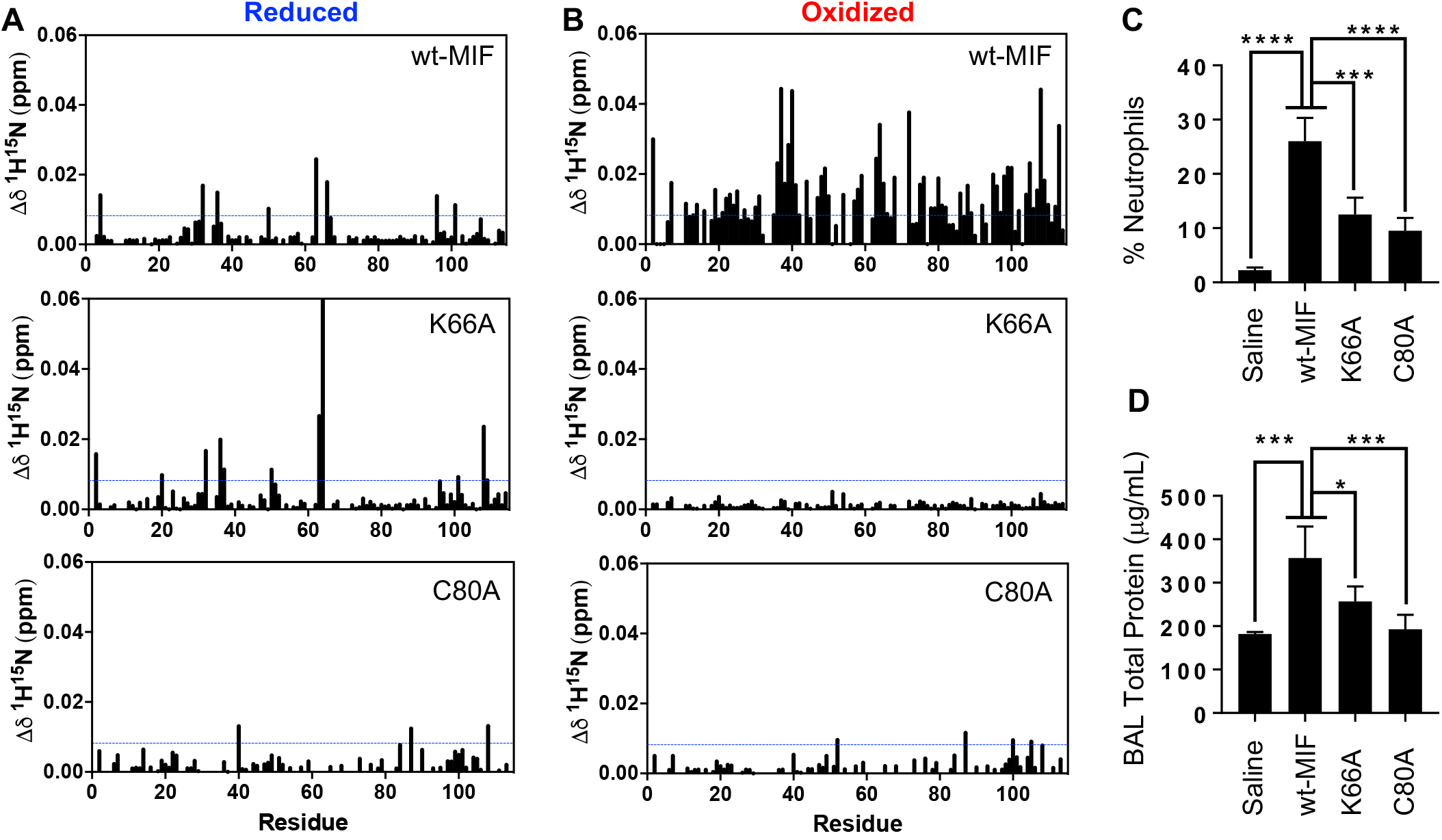
Mutation of the redox-sensitive K66 and C80 residues alters the redox-dependent MIF structure and hinders the ability of MIF to activate CD74 *in vivo*. K66A and C80A mutations alter the combined ^1^H-^15^N chemical shift profiles of reduced **(A)** and oxidized **(B)** MIF, relative to redox-neutral K66A and C80A samples used as references. Blue lines denote 1.5σ above the 10% trimmed mean of all shifts. Combined ^1^H-^15^N chemical shift plots for reduced and oxidized wt-MIF are repeated here from Figure 1 for comparison. **(C)** K66A and C80A mutations significantly decreased CD74-dependent neutrophil recruitment in murine lungs *in vivo*. **(D)** Attenuated CD74-dependent neutrophil recruitment by K66A and C80A also led to significantly decreased pulmonary edema *in vivo*. (n = 4 in each group, *p<0.05, ***p≤ 0.001 and ****p≤0.0001). Data are expressed as mean ± SEM.

To understand whether this altered redox behavior foreshadowed changes in MIF function, we evaluated the K66A and C80A variants in neutrophil recruitment assays in murine lungs, where we measured total immune cells, percentage of neutrophils in BAL fluid, and protein content (as a marker of alveolar capillary leak/pulmonary edema). This *in vivo* assay quantifies neutrophil recruitment on the surface of alveolar macrophages, an established metric for the MIF-induced activation of the pro-inflammatory CD74 receptor.^34^ The assay is indirect, using chemokines released by activated macrophages to stimulate CXCR2 receptors on neutrophils (that do not directly express CD74) to induce their migration to murine lungs.^35^ We found that wt-MIF significantly increased the total BAL fluid cell counts over saline controls, while the K66A and C80A variants showed a reduced number of total BAL fluid immune cells (**Figure S11**). The neutrophil influx and pulmonary edema protein levels were significantly increased in the wt-MIF group relative to vehicle controls (**Figures 5C, 5D**). In contrast, we observed significant decreases in the groups to which the MIF variants were delivered, as compared to wt-MIF. Together, our data indicate that the K66A and C80A variants have a functional effect in reducing the inflammatory response of MIF in the murine lung. These findings also suggest that structural and/or dynamic sensitivity of residues within MIF can provide a novel route for achieving functional control beyond the known MIF active sites.

## Discussion

We used solution redox potentials to modulate the MIF structure and dynamics in order to reveal changes in conformational equilibria and redox-sensitive residues that may be latent functional sites within its structure. This work builds upon prior studies implying redox sensitivity of MIF^11,22,36,37^ as well as our recent paper highlighting a series of allosteric residues linking the enzymatic (tautomerase) and CD74 binding sites of MIF.^38^ Here, we demonstrate that examination of MIF redox sensitivity is another useful avenue for probing function beyond its established active sites. Since MIF participates in both intracellular and extracellular interactions,^39^ where the redox environments differ, understanding the implications of its redox dependence may also inform its pro-inflammatory mechanisms and functional promiscuity, which has been investigated at the biochemical, but not the structural, level. A previous characterization of the association of MIF with p53 tumor suppressor suggested the interaction to be redox-dependent, having shown a significant decrease in coprecipitation in the presence of a reductant.^40^ Further, interaction of MIF with ribosomal protein S-19 (RPS-19) was shown to occur more strongly under reducing conditions.^41^ In addition, MIF is known to be post-translationally modified by glycation in the brain tissue of Alzheimer’s patients. Most notably, an altered electrophoretic migratory behavior of glycated MIF was observed and demonstrated *in vitro* by incubating MIF with glucose under oxidizing conditions.^42^

Redox signatures of inflammatory diseases, most of which implicate MIF, can also be markedly different from those of healthy humans. The redox potential of epithelial lining fluid is - 110 to -40 mV in the lungs of asthmatics, which is significantly more oxidizing than that of healthy humans (−150 to -200 mV).^27^ Further, the synovial fluid of rheumatoid arthritis patients has been shown to be highly oxidizing relative to that of a non-arthritic control group.^43,44^ A significant body of literature suggests that when overexpressed at sites of inflammation,^16^ MIF can endure large swings in environmental redox potential, consistent with our determination of the MIF redox potential *in vitro* using solution NMR^45,46^ (**Figure S12**, midpoint 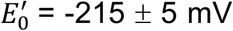, near normal physiological range). NMR spin relaxation studies show redox-dependent modulation of local flexibility in MIF, with motion observed on multiple timescales. oxMIF, in particular, exhibits the most significant structural and dynamic changes, with widespread NMR spectral perturbations and CPMG relaxation dispersion that show oxidizing potentials stimulate flexibility in ∼34% of the MIF structure, including its redox-active residues. However, the majority of these flexible regions have not been confirmed to harbor any functional impact in prior work.

It remains unclear how this relatively compact (12.5 kDa monomer) and symmetrical homotrimer accommodates a wide range of non-overlapping activities. A 2018 study showed that nanosecond dynamics were important for organization of the MIF-CD74 binding interface,^6^ and studies of a MIF epitope containing its ^56^CALC^59^ motif showed selective binding to the Fab fragment of the BaxB01 antibody under oxidizing conditions, after rearrangement of the Cys sulfhydryls.^22^ Oxidizing conditions appear to promote a more flexible solvent channel, which we recently showed to be central to an allosteric mechanism regulating tautomerase and CD74 activation involving motion on multiple timescales.^38^ Given the number of MIF functions that have yet to be explored at the molecular level and the fact that ∼100 X-ray crystal structures of MIF mutants, substrate-, and inhibitor-bound complexes have negligible variation in static structure but vastly different NMR spectra, it is plausible, if not likely, that additional regions of functional importance exist in MIF.

Here, we asked if changes in solution redox potential could reveal some of these latent sites by modulating the structural and/or dynamic signatures of MIF. We experimentally altered the redox potential of wt-MIF NMR samples, enabling us to identify redox-dependent structural changes in residues that may be important for function. We mutated two such residues, K66 and C80, that showed redox-sensitivity in wt-MIF and observed structural effects on MIF redox behavior as well as altered functional responses. Having further confirmed the propensity of C80 for redox-driven modification by mass spectrometry, we speculate that MIF could act as a sensor that toggles its conformation depending on its cellular environment to engage with varied binding partners, as was previously suggested in a conformational control scheme for the ^56^CALC^59^ motif. The more extensive but partially-overlapping set of residues affected by the redox environment suggests the regulatory network controlling non-overlapping MIF functions may be larger than previously reported, where additional “allosteric nodes” may be revealed only when MIF is exposed to stimuli, such as environmental factors associated with inflammation, serving as a natural means of activating latent residues to expand MIF functions under altered cellular conditions. Speculating further, targeting small molecules to redox sensitive residues may prevent MIF from engaging in certain downstream cascades. Future studies of protein-protein interactions with supposed binding partners of MIF, such as thioredoxin^47,48^ or ribosomal protein S19,^41^ as well as with small molecules, can evaluate this mechanism. Another interesting avenue warranting further exploration is the redox-dependent interaction of MIF with its known inhibitors or with novel compounds, as it presents an opportunity to preferentially target oxMIF, a form of the protein that may appear predominately within the inflammatory state.

## Materials and Methods

### Protein expression, purification, and redox sample preparation

Wild-type (wt) and/or human MIF variants cloned into a pET11b vector were grown in lysogeny broth (LB) for biochemical studies and in isotopically enriched M9 minimal medium supplemented with CaCl_2_, MgSO_4_, and MEM vitamins for NMR, after adapting BL21(DE3) cells to D_2_O. Small cultures of MIF were grown overnight in LB at 37 °C and used to inoculate cultures containing 50% D_2_O the following morning, which were grown 8 – 10 hours at 37 °C and then used to inoculate cultures containing 95% D_2_O, which were incubated at 37 °C for 12 hours. The cells were collected by centrifugation and resuspended in M9 medium (1 L, 100% D_2_O) supplemented with ^15^NH_4_Cl (Cambridge Isotope Laboratories) and ^12^C_6_H_12_O_6_ as the sole nitrogen and carbon sources, respectively. Cultures were grown to an OD_600_ of 0.8 – 1.0 and induced with 1 mM isopropyl β-D-1-thiogalactopyranoside (IPTG). Following 16-18 hours of post-induction growth, the cells were harvested by centrifugation.

The cells were resuspended in ∼30 mL of an ice-cold buffer of 20 mM Tris-HCl, 20 mM NaCl, and 1 mM EDTA at pH 7.4 supplemented with protease inhibitors, lysed by ultrasonication, clarified by centrifugation, filtered, and purified with a 120 mL Macro-Prep High Q column(BioRad) pre-equilibrated with the lysis buffer. MIF does not bind the Q resin and is found in the flow-through. MIF was purified to its final form on a HiLoad 26/600 Superdex 200 size exclusion column (GE Healthcare). For NMR samples, MIF was dialyzed into a buffer containing 20 mM sodium phosphate, 1 mM EDTA, and 7.5% D_2_O at pH 7.0. Final concentrations of MIF were 0.5 – 1 mM determined using ε_280_ = 12,950 M^-1^ cm^-1^.^49^

Oxidation of MIF was achieved by addition of H_2_O_2_ to a final concentration of 6 mM in an aliquot of 0.5 – 1 mM MIF. Reduced MIF was prepared with a 6 mM final concentration of DTT. Redox-neutral samples were prepared with equimolar amounts of DTT/H_2_O_2_. Stocks of the oxidizing and reducing agents were freshly prepared in a degassed buffer of 20 mM sodium phosphate, 1 mM EDTA, and 7.5% D_2_O at pH 7.0. Following addition of oxidant or reductant to MIF samples, the NMR tubes were briefly degassed with nitrogen and sealed with Parafilm (Bemis). The samples were equilibrated in the presence of redox reagents for at least six hours. *Circular Dichroism (CD) Spectroscopy*

CD spectra and thermal unfolding experiments were recorded on a JASCO J-815 spectropolarimeter equipped with a variable temperature Peltier device. Denaturation curves of 10 μM MIF were recorded at 218 nm in a 2 mm quartz cuvette. The temperature range for each scan was 20 – 100 °C (293 K – 373 K). Thermodynamic parameters were extracted via nonlinear curve fitting of CD data to **Equation 1** in GraphPad Prism:

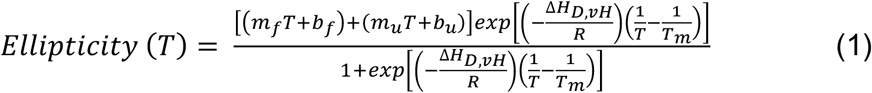

where *m*_f_, *b*_f_, and *m*_u_, *b*_u_ are the slopes and y-intercepts of the folded (low temperature) and unfolded (high temperature) regions of the melting curve, *R* is the gas constant, *T*_m_ and Δ*H*_D,vH_ are the unfolding midpoint and van’t Hoff enthalpy of denaturation at *T*_m_, respectively. Free energy analysis was performed as described elsewhere,^50^ using values of Δ*C*_p_, the unfolding heat capacity, estimated from the report of Privalov.^51,52^ The differences in apparent unfolding enthalpies, entropies, and free energies at a reference temperature (*T*_REF_) of 298 K were calculated as;

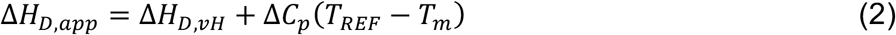

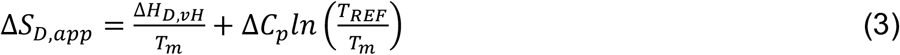

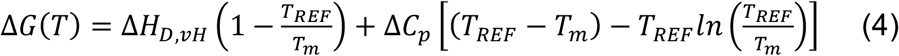

These data are reported as “apparent” values due to the ∼90% reversibility of the MIF unfolding transition.

### NMR Spectroscopy

NMR experiments were performed on a Bruker Avance NEO 600 MHz or Bruker Avance III HD 850 MHz spectrometer at 30 °C. NMR data were processed using NMRPipe^53^ and analyzed in Sparky^54^ along with in-house scripts. The pH of NMR samples was monitored after addition of redox reagents to ensure that NMR spectral changes were due to reduction/oxidation. NMR assignments for wt-MIF and the K66A and C80A variants were confirmed from standard triple resonance experiments. Combined chemical shift perturbations (Δδ) were determined by 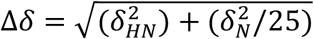. The HSQC of oxidized MIF was processed with an exponential window function in both the direct and indirect dimensions in NMRPipe. The peak intensity and volume for each pair of slow exchanging peaks were calculated in SPARKY, where peak volume was calculated by integrating each peak as a Lorentzian. The minor state populations were estimated by taking the volume of the minor peak and dividing by the total volume of the slow exchanging pairs.

NMR spin relaxation experiments were performed using TROSY-based pulse sequences adapted from Palmer and coworkers.^55^ Relaxation data were acquired with the ^1^H and ^15^N carriers set to the water resonance and 120 ppm, respectively. Longitudinal relaxation rates (*R*_*1*_) were measured with *T*_1_ delays of 0, 20, 60, 100, 200, 600, 800, and 1200 ms. Transverse relaxation rates (*R*_*2*_) were collected with 1.0 ms spacing between 180° CPMG pulses at total relaxation delays of 0, 1, 2, 4, 8, and 10, and 12 ms. The recycle delay in these experiments was 2.5 s. Longitudinal and transverse relaxation rates were extracted by non-linear least squares fitting of the peak heights (major peaks in cases of slow exchange) to a single exponential decay using in-house software. Uncertainties in these rates were determined from replicate spectra. The heteronuclear cross-relaxation rate (^1^H-[^15^N] NOE) was obtained by interleaving pulse sequences with and without proton saturation and calculated from the ratio of peak heights from these experiments. Carr-Purcell-Meiboom-Gill (CPMG) NMR experiments were performed at 30 °C with a constant relaxation period of 40 ms, a 2.0 s recycle delay, and τ_cp_ points of 0, 25, 50, 75, 100, 150, 250, 500, 750, 800, 900, and 1000 ms. Relaxation dispersion profiles were generated by plotting *R*_2_ vs. 1/τ_cp_ and exchange parameters were obtained from fits of these data carried out with in-house scripts. Uncertainties were obtained from replicate spectra.

NMR spin-relaxation rates were fit to one of five semi-empirical forms of the spectral density function using model-free analysis.^56,57^ Fitting of motional parameters was performed in RELAX.^58,59^ The criteria for inclusion of residues in the diffusion tensor estimate relied on the method of Tjandra and coworkers.^47^ N-H bond lengths were assumed to be 1.02 Å and the ^15^N chemical shift anisotropy was set to -160 ppm. During the model selection process, the diffusion tensor parameters were optimized simultaneously, and model selection was repeated until the optimized tensor parameters and order parameter (S^2^) did not differ from those of the previous iteration.

### Mass Spectrometry

Cysteine and cystine residues of purified MIF equilibrated under each of the redox conditions were alkylated in a stepwise process with iodoacetamide (IAA) followed by *N*-ethylmaleimide (NEM). Parallel reaction monitoring-mass spectrometry captured the oxidation states of C80 under different redox conditions. Skyline software^60^ was used to calculate the relative abundance of cysteine and cystine at the C80 position, indicated by binding to IAA or NEM. Additional methodological details can be found in the **Supporting Information**.

### In Vivo Neutrophil Recruitment Assay

All animal studies were approved by the Institutional Animal Care and Use Committee of Cooper University Healthcare, Camden, NJ. The wt mice of genetic background strain (C57BL6/J) were purchased from the Jackson Laboratory (Bar Harbor, ME). Mice were housed at the pathogen-free animal facility at Cooper University Healthcare. All experiments were done in 9-10 week old male mice. Mice were administered a one-time intratracheal instillation of 100 µl of normal saline alone (vehicle) or normal saline solution containing 1 µg of wt-MIF, K66A MIF or C80A MIF. These mice were sacrificed after 6 h of vehicle only or vehicle + experimental agent administration. Bronchoalveolar lavage (BAL) was performed by cannulating the trachea with a blunt 22-gauge needle and lavaging both lungs with 800 µl of sterile PBS solution. Bronchoalveolar lavage fluid (BALF) was harvested, and total cell counts in BALF were determined using the TC20 automated cell counter (Bio-Rad Laboratories, Inc., Hercules, CA). The differential cell counts were performed on cells cytocentrifuged onto glass slides (Fisher Scientific) stained with the Hema 3 Staining System (Fisher Diagnostics, Middletown, VA) and cell differential was tabulated using light microscopy. Total protein concentration in the BAL fluid was measured using the Pierce™ BCA assay kit (Thermo Scientific, Rockford, IL), as previously described.^6^

## Acknowledgments

The authors thank J. Patrick Loria for helpful discussions. This work was supported by Rhode Island Foundation Grant GR5290658 and funds from the Office of the Vice President for Research at Brown University (to GPL).

## Competing Interests

The authors declare no competing interests.

## Notes

### Competing Interest Statement

The authors have declared no competing interest.

