## Supplementary Data for "Redox-dependent Structure and Dynamics of Macrophage Migration Inhibitory Factor Reveal Sites of Latent Allostery"

**Supplemental Mass Spectrometry Methods**

**Figure S1.** Redox-dependent secondary structure and stability of MIF

**Figure S2:** NMR spectra of oxidized and reduced MIF

**Figure S3:** Analysis of slow exchange peaks observed in oxidized MIF

**Figure S4:** Summary of redox-dependent modification of MIF

**Figure S5:** $R_1$, $R_2$, $^1$H-[$^{15}$N] NOE spin relaxation data for redox-modulated MIF

**Figure S6:** Representative CPMG relaxation dispersion data

**Figure S7:** Raw mass spectrometry data

**Figure S8:** C80 reactivity and conformational landscape

**Figure S9:** Redox-dependent NMR spectra of K66A and C80A MIF variants

**Figure S10**: Figure S10: Structural stability of K66A and C80A MIF variants

**Figure S11:** Diminished *in vivo* activity of K66A and C80A MIF

**Figure S12:** Determination of MIF redox potential from NMR chemical shifts

### Supplemental Mass Spectrometry Methods

**Proteomics.** Purified samples of MIF equilibrated under oxidizing and redox-neutral conditions were incubated with a final concentration of 15 mM iodoacetamide (IAA, room temperature, 1 hour in the dark) to alkylate any reduced cysteine residues. Subsequently, cysteines that were in an oxidized state were reduced with a final concentration of 10 mM DTT (room temperature, 30 minutes). Prior to alkylation with *N*-ethylmaleimide (NEM), the sample solution was adjusted to pH 6.5 with 1% formic acid to minimize over-labeling of lysine and N-termini.[1] These samples were then incubated with a final concentration of 15 mM NEM (room temperature, 1 hour in the dark). The reaction was quenched by the addition of DTT to a final concentration of 10 mM. Purified MIF samples equilibrated under reducing conditions were treated with only IAA or only NEM at 15 mM (room temperature, 1 hour in the dark) and then quenched with 10 mM DTT. Labeled MIF samples were dried using vacuum centrifugation, resuspended in 200 mM EPPS, pH 8.5, and digested overnight at 37° C with Trypsin/Lys-C protease mixture at a 50:1 protein to protease ratio. Digested samples were again dried and resuspended in a 5/1 (v/v) solution of acetonitrile and formic acid prepared in ultrapure water. The samples were subjected to C18 solid-phase extraction with columns prepared in-house. The final samples were dried and reconstituted in a solution of 5/2 (v/v) of acetonitrile and formic acid in ultrapure water to a final concentration of 12.5 ng/μL for liquid chromatography-mass spectrometry (LC-MS) analysis.

**Liquid Chromatography and Parallel-Reaction Monitoring (PRM) Mass Spectrometry.** Mass spectrometric data was collected on an Orbitrap Fusion Eclipse (ThermoFisher Scientific, San Jose, CA) coupled to a FAIMSPro (ThermoFisher Scientific, San Jose, CA) and a Proxeon Easy-nLC 1200 (ThermoFisher Scientific, San Jose, CA). Peptides were separated on a 100 μm inner diameter microcapillary column packed with ~ 35 cm of Reprosil-Pur 120 C18 resin (3 μm, AQ, ESI Source Solutions, Woburn, MA). For each analysis, 12.5 ng of MIF sample was loaded onto the column and separated using a 30 min linear gradient from 100% A (5% ACN, 0.125% formic acid) to 50% B (59% ACN, 0.125% formic acid). PRM analyses used an inclusion list of 23 targets generated from Skyline software that included precursor mass-to-charge ratio (*m/z*) and precursor charge state for alkylated and unmodified MIF peptides. All targeting was unscheduled and occurred throughout the duration of the experiment. The scan sequence began with an MS1 spectrum (Orbitrap resolution 120 000) followed by targeted MS2 scans of peaks corresponding to precursors in the target list. Precursors were isolated by the quadrupole mass analyzer with a 1.2 *m/z* wide window and fragmented using higher-energy collision dissociation (HCD). Fragments were detected in the ion trap using the rapid scan rate and a maximum injection time of 25 ms. The above scans were collected using alternating FAIMS compensation voltages of -40 and -60 V and a dispersion voltage of -5000 V.

**Data Analysis.** All raw files were imported into the Skyline software package (Version 20.2.0.343)[2] without modification. Peak areas corresponding to peptide 78-LLCGLLAER-86 with +57 Da (IAA) or + 125 Da (NEM) modification at the C80 position were determined using the built-in peak-boundary adjustment tool. Relative amounts of C80 oxidation were determined against the intensity of all peaks in the chromatogram for each condition. Five replicates from two sample preparations were analyzed for experimental variance.

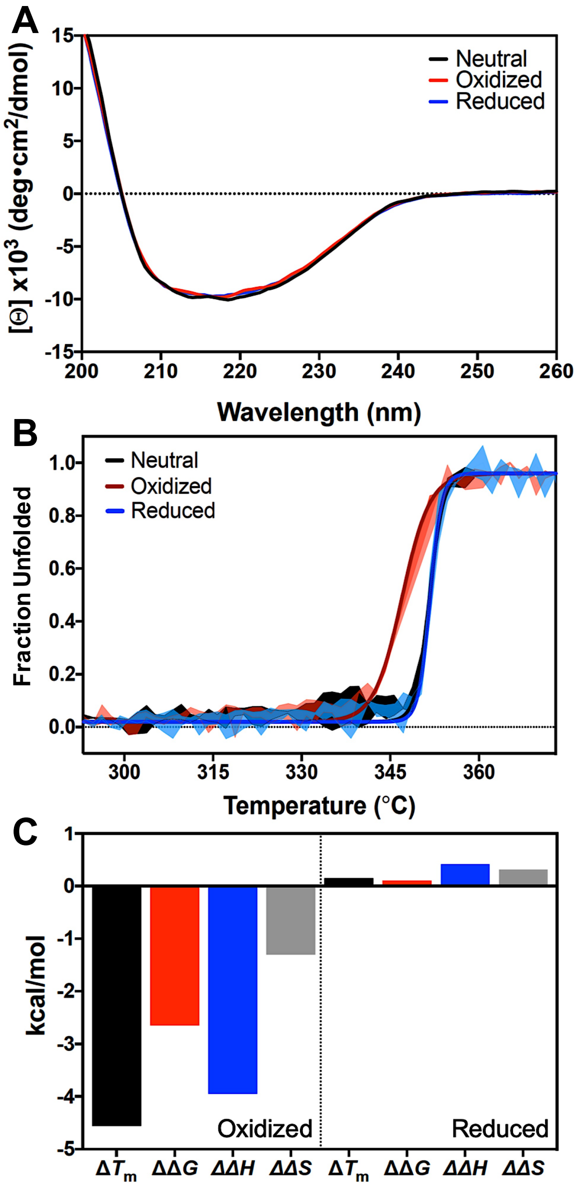


**Figure S1.** Redox-dependent secondary structure and stability of MIF. **(A)** Far-UV circular dichroism spectra of redox-neutral (black), oxidized (red) and reduced (blue) MIF. **(B)** Redox-dependent thermal denaturation profiles of MIF ($\lambda$ = 218 nm) normalized as a fraction of unfolded protein. Solid lines represent fits of the data to **Equation 1**, and shaded areas represent the error in the data points over multiple unfolding experiments. **(C)** Summaries of apparent redox-dependent changes in $T_m$ (black), free energy (red), enthalpy (blue), and entropy (gray) of MIF. These data are reported per trimer using $\Delta C_p$ = 4.8 kcal/molK (114 residues per monomer, 342 residues total) and a reference temperature of 351.7 K, the $T_m$ of redox-neutral MIF (thus, $\Delta\Delta G_{app}$ for redox-neutral MIF = 0).

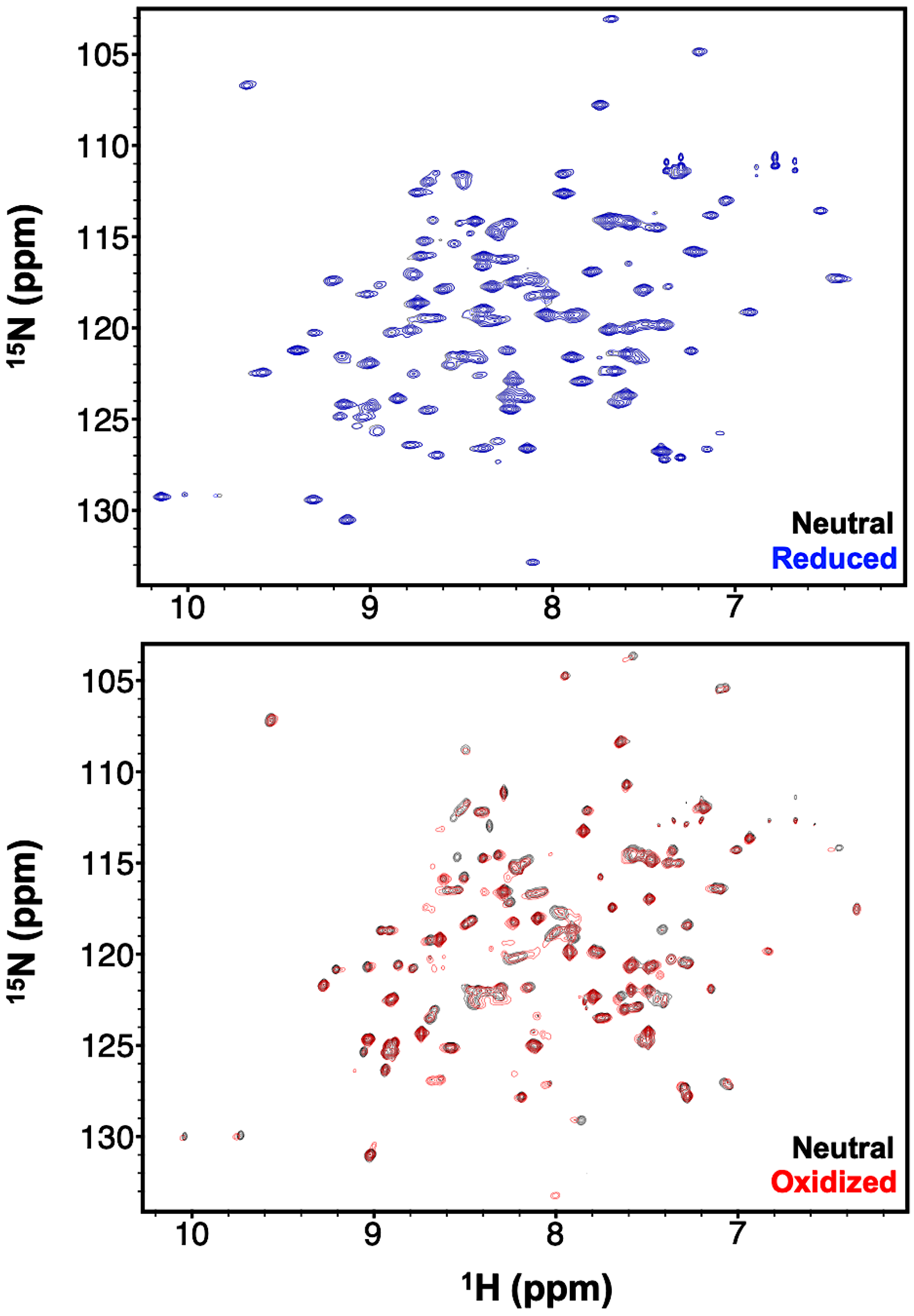


**Figure S2:** $^{1}H$-$^{15}N$ TROSY HSQC NMR spectra of MIF collected under reduced (blue) and oxidized (red) solution conditions. Spectra are overlaid with that of redox-neutral MIF (black).

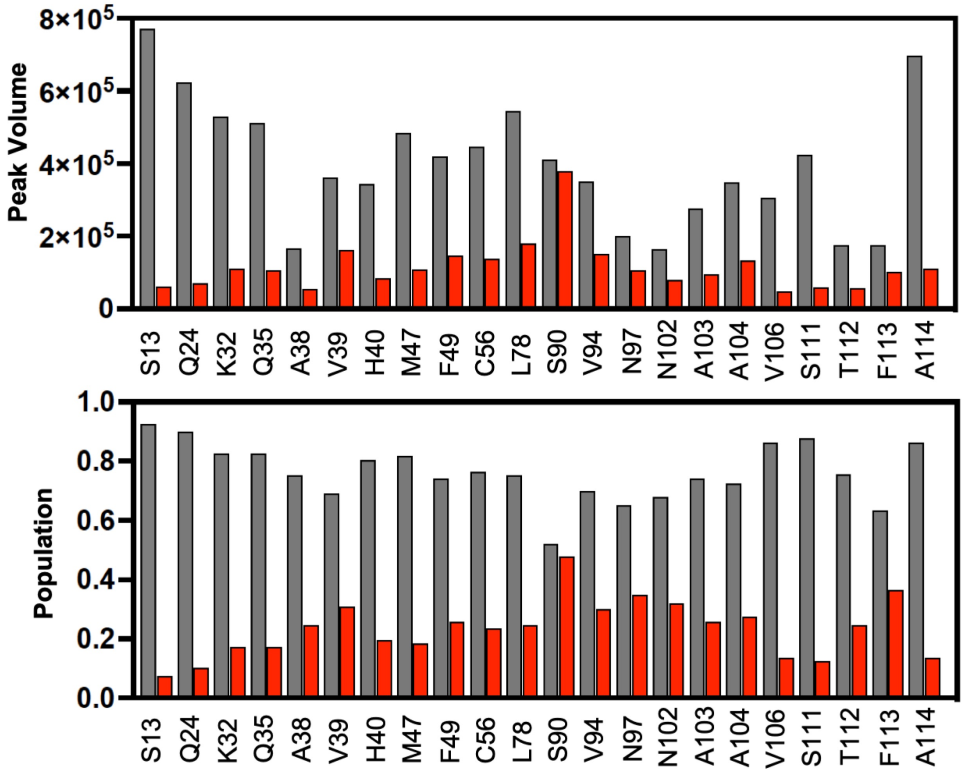


**Figure S3:** Peak volumes and populations of the major (gray) and minor (red) peaks at sites of slow exchange in oxidized MIF. The peak intensity and volume for each pair of slow exchanging peaks were calculated in SPARKY, where peak volume was calculated by integrating each peak as a Lorentzian. The minor state populations were estimated by taking the volume of the minor peak and dividing by the total volume of the slow exchanging pairs.

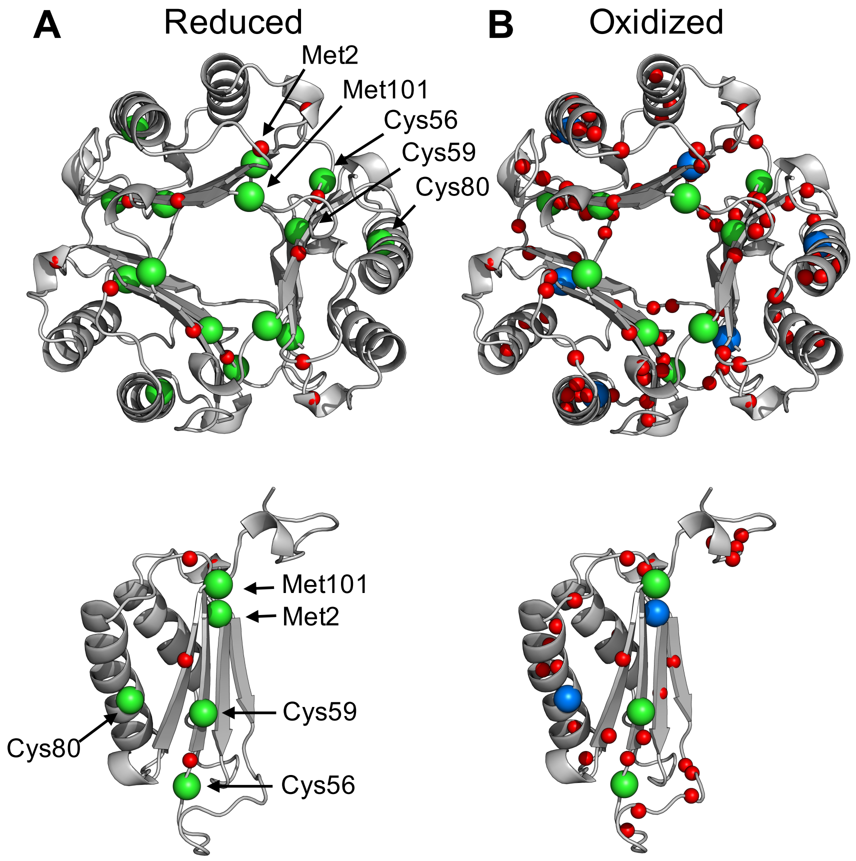


**Figure S4:** Sites of redox-sensitive chemical shift perturbations caused by reducing **(A)** and oxidizing **(B)** conditions are mapped onto the MIF structure (PDB: 1MIF). In both cases, spheres represent spectral perturbations relative to redox-neutral MIF. Traditionally redox-sensitive Cys and Met residues are shown in larger green spheres and chemical shift perturbations are shown in smaller red spheres. Sites where chemical shift perturbations occur at Cys/Met residues are shown in larger blue spheres.

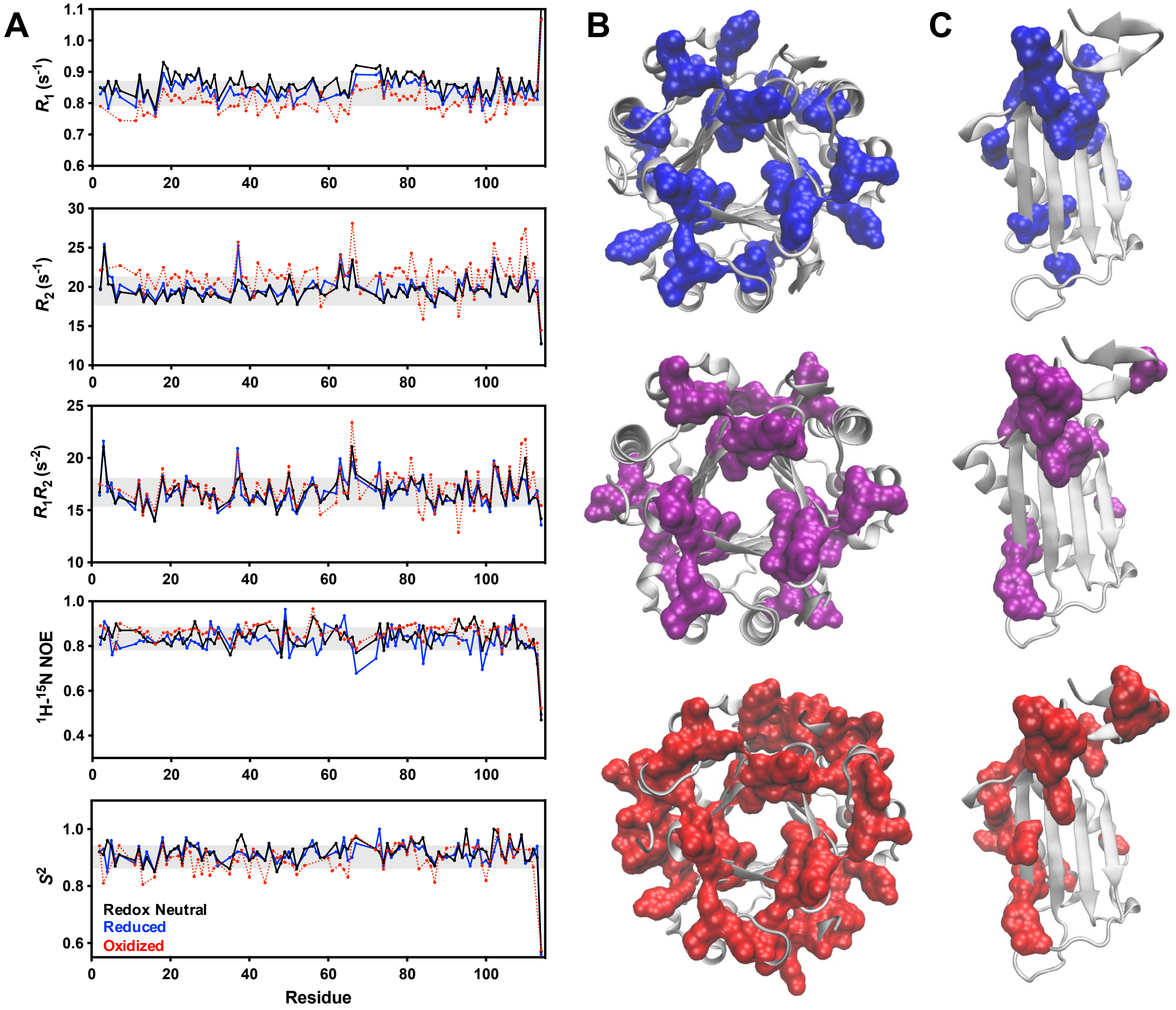


**Figure S5. (A)** Plots of $R_1$, $R_2$, $R_1R_2$, and the $^1$H-[$^{15}$N] NOE and order parameter ($S^2$) for reduced (blue), redox neutral (black), and oxidized (red dash) MIF. Gray shaded bars denoted $\pm 1.5\sigma$ of the 10% trimmed mean of all data collected for each parameter. Data from the $R_1R_2$ and $S^2$ plots in **(A)** are the basis of the correlation profiles shown in main text **Figure 3**. Sites suggestive of μs – ms chemical exchange in MIF determined from $R_1R_2$ values $1.5\sigma$ above the 10% trimmed mean of the data are mapped onto the MIF trimer **(B)** and monomer **(C)** structures (PDB: 1MIF) as a function of solution redox potential. Surfaces correspond to reduced (blue), redox-neutral (purple), and oxidized (red) MIF.

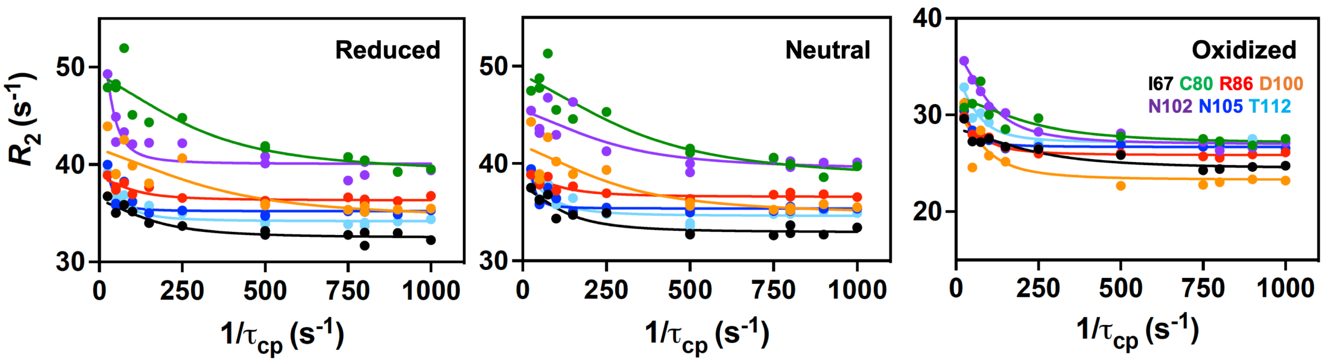


**Figure S6.** Representative redox-dependent CPMG relaxation dispersion profiles for residues in MIF. These data highlight selected examples of conserved ms timescale motions in MIF across redox regimes, while variations in dynamic profiles are discussed in greater detail in main text **Figure 3**.

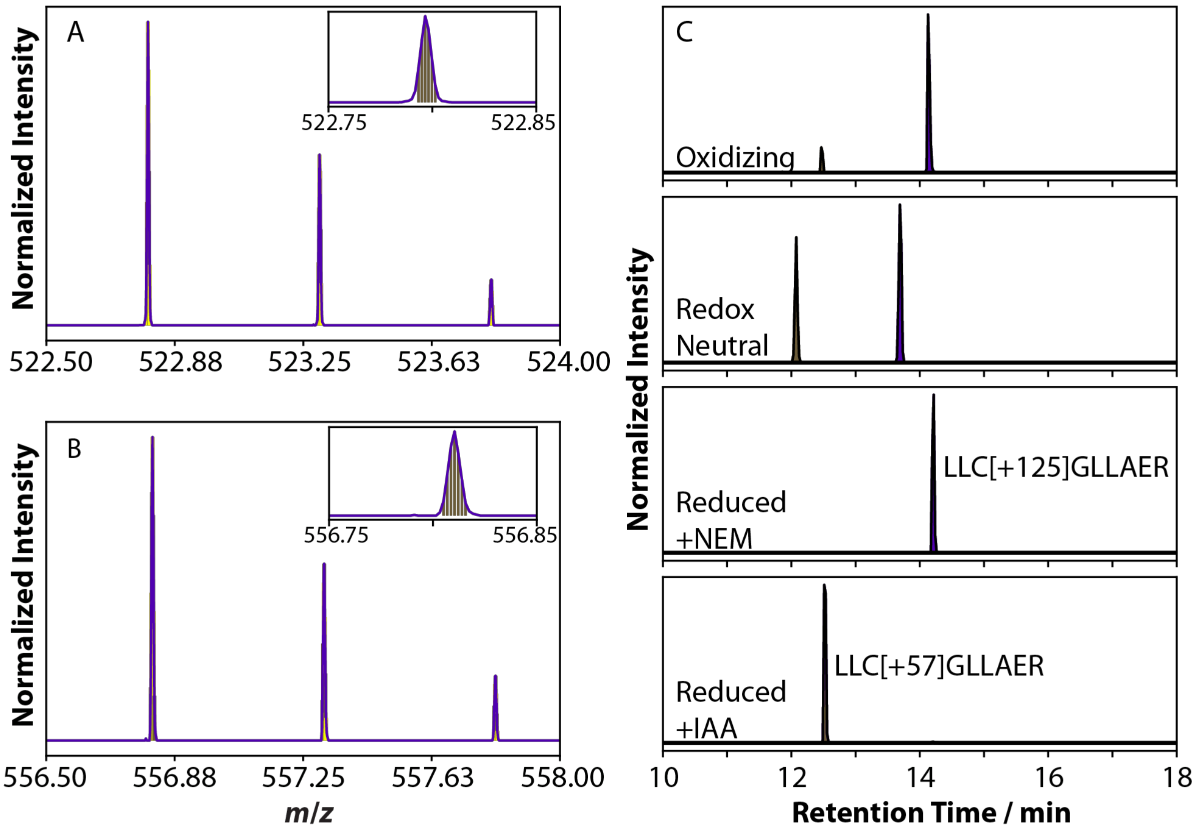


**Figure S7.** Raw mass spectrometry data processed using Skyline software. Agreement between raw spectra and Skyline-predicted spectra of C80-containing peptide precursor modified by IAA **(A)** and NEM **(B)**. The separation of the differentially modified C80 in the LC trace is shown in **(C)** with the two experimental conditions shown in the top two traces and the reduced-control traces shown in the bottom two plots.

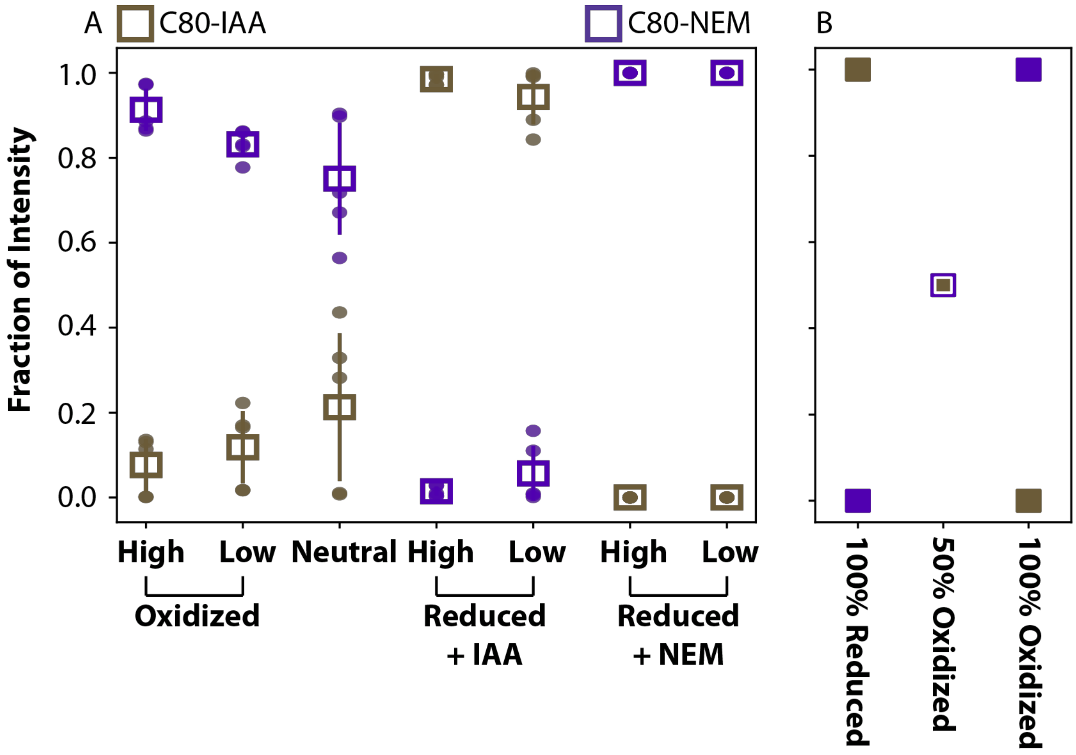


**Figure S8.** C80 reactivity and conformational landscape depends on the concentration of oxidant in the environment. As the concentration of oxidizing species is increased, the amount of C80 in cystine form increases relative to lower oxidant concentrations **(A)**. In **(B)**, the expected contributions to the total ion intensity in the chromatogram for fully reduced, 50% oxidizing, and fully oxidized C80 is shown.

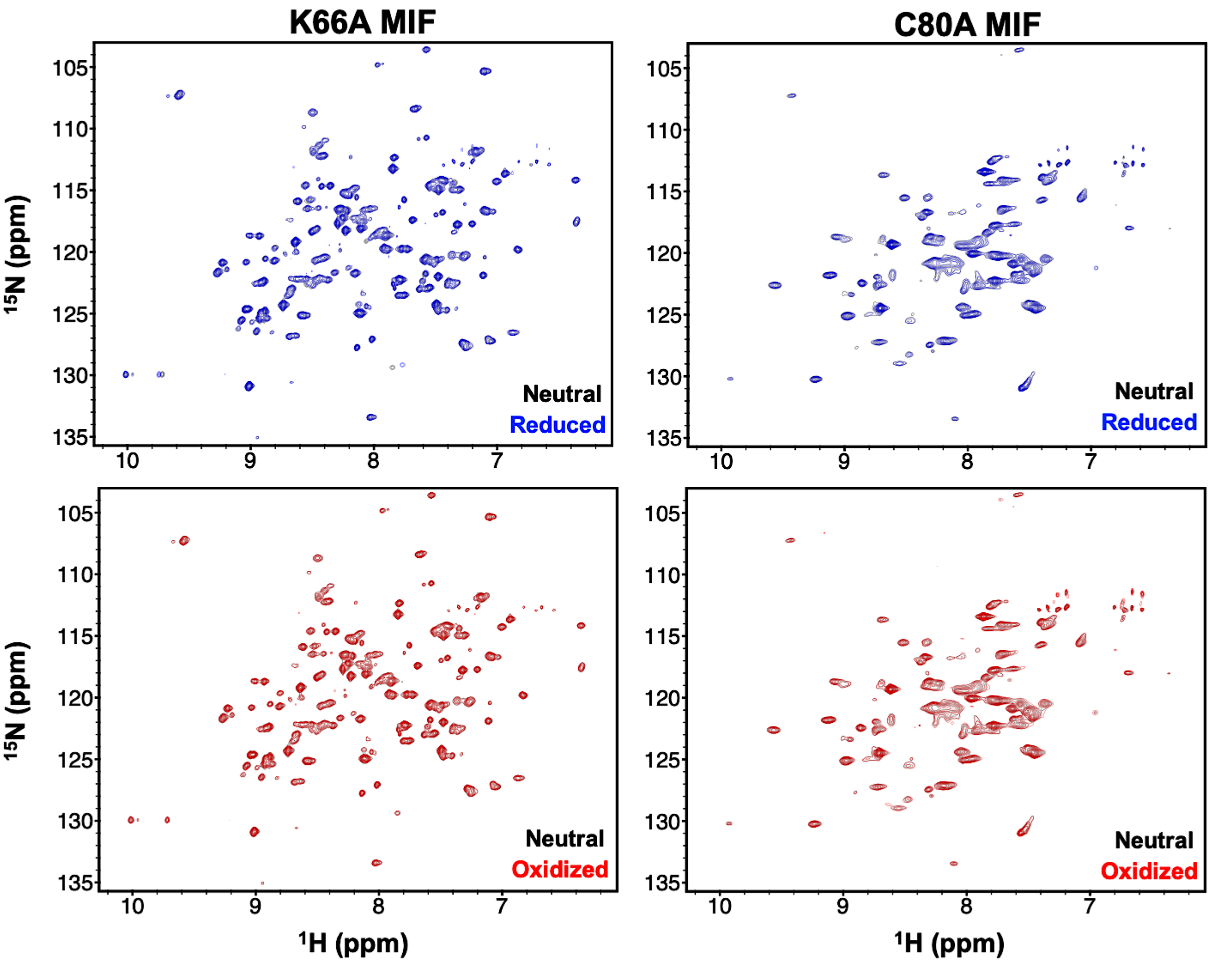


**Figure S9.** $^{1}$H-$^{15}$N TROSY HSQC NMR spectra of K66A **(A)** and C80A **(B)** MIF collected under reduced (blue) and oxidized (red) solution conditions. Spectra are overlaid with that of redox-neutral MIF variants (black). The high degree of line broadening in C80A spectra occurs upon mutation but does not change as the solution redox potential is altered. These data are the basis for the chemical shift perturbations displayed in main text **Figure 5**.

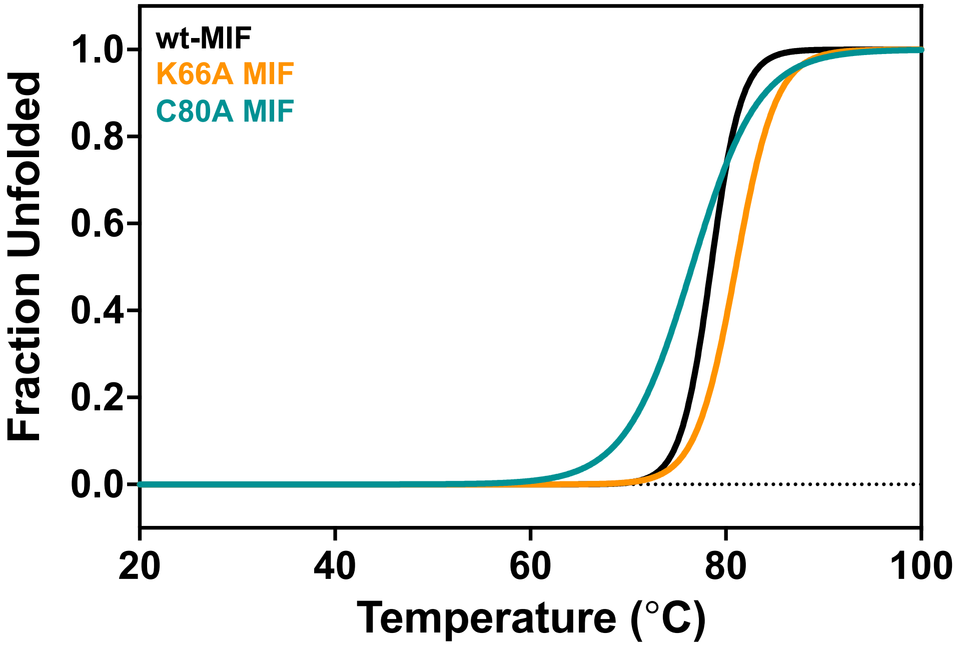


**Figure S10:** Structural stability of K66A and C80A MIF variants. Thermal denaturation profiles ($\lambda$ = 218 nm) of K66A MIF (orange), C80A MIF (teal), and wt-MIF (black) are normalized as a fraction of unfolded protein. Despite minor stability changes resulting from single point mutations in the MIF variants, as compared to wt-MIF, K66A and C80A MIF are fully folded under NMR experimental conditions (30 °C), which is consistent with observed stability changes of other MIF variants as previously reported.[3]

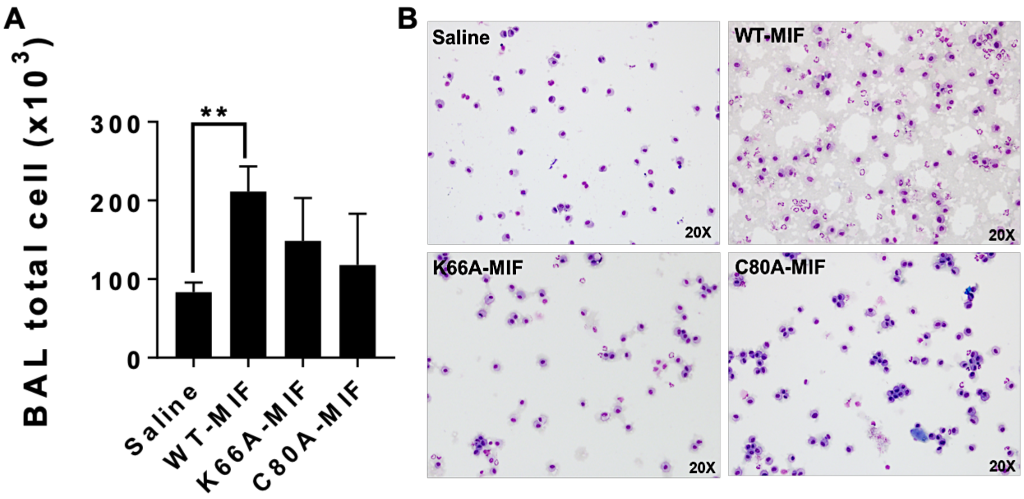


**Figure S11. (A)** Mutations at K66 and C80 decrease total immune cell recruitment in lungs *in vivo*. **(B)** Representative images of Hema staining in BAL fluid cell pellets show decreased neutrophil recruitment in murine lungs by K66A and C80A variants as compared to wt-MIF administered as a one-time intra-tracheal dose. (n = 4 in each group, **p ≤ 0.01). Data are expressed as mean ± SEM.

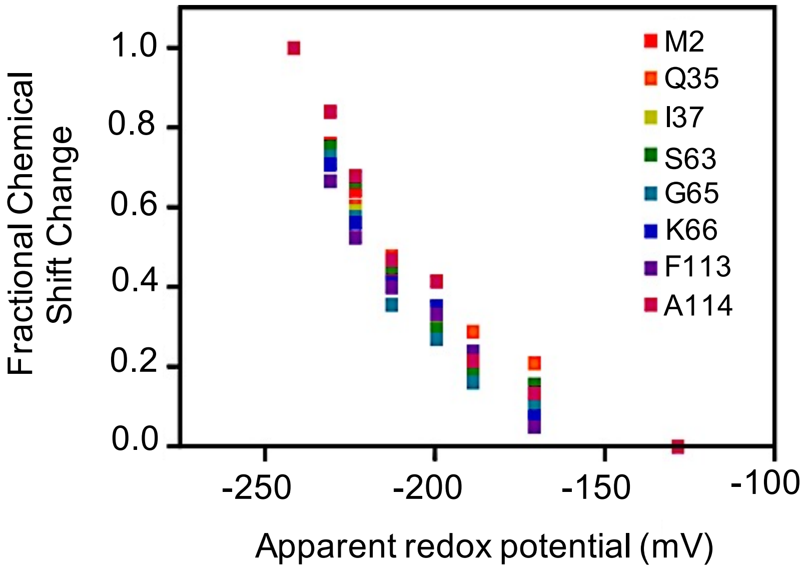


**Figure S12.** Determination of the MIF redox potential by adapting the method of Freund and coworkers.[4] A redox-neutral sample of $^{15}N$ MIF (0.5 mM) was titrated with reduced glutathione (GSH) to achieve [GSH] = 1, 2, 3, 5, 7.5, 10, and 15 mM. $^{1}H$-$^{15}N$ HSQC NMR spectra were collected at each GSH concentration and combined chemical shift values for residues with measurable responses to GSH (a selection is shown here) were normalized against the total perturbation for that residue over the entire titration. The MIF redox potential was calculated from the GSH:GSSG ratio via the Nernst equation $E_0' = E_{GSH}' - (RT/nF)\ln(K_{eq})$, where the midpoint is equal to the redox potential.
